# A conserved viral amphipathic helix governs the replication site-specific membrane association

**DOI:** 10.1101/2021.12.09.472036

**Authors:** Preethi Sathanantham, Wenhao Zhao, Guijuan He, Austin Murray, Emma Fenech, Arturo Diaz, Maya Schuldiner, Xiaofeng Wang

**Affiliations:** School of Plant and Environmental Sciences, Virginia Tech, Blacksburg, Virginia 24061, United States; Institute of Plant Protection, Jiangsu Academy of Agricultural Sciences, Key Lab of Food Quality and Safety of Jiangsu Province-State Key Laboratory Breeding Base, Nanjing 210014, China; Fujian Province Key Laboratory of Plant Virology, Institute of Plant Virology, Fujian Agriculture and Forestry University, Fuzhou, Fujian 350002, China; Department of Molecular Genetics, Weizmann Institute of Sciences, Rehovot 7610001, Israel; Department of Biology, La Sierra University, Riverside, California 92505, United States

**Author notes:** Correspondence and requests for materials should be addressed to XW.

**Keywords:** amphipathic helix, membrane binding, viral replication complexes

## Abstract

Positive-strand RNA viruses assemble their viral replication complexes (VRCs) on specific host organelle membranes, yet it is unclear how viral replication proteins recognize and what motifs or domains in viral replication proteins determine their localizations. We show here that an amphipathic helix, helix B in replication protein 1a of brome mosaic virus (BMV), is necessary for 1a’s localization to the nuclear endoplasmic reticulum (ER) membrane where BMV assembles its VRCs. Helix B is also sufficient to target soluble proteins to the nuclear ER membrane in yeast and plant cells. We further show that an equivalent helix in several plant- and human-infecting viruses of the alphavirus-like superfamily targets fluorescent proteins to the organelle membranes where they form their VRCs, including ER, vacuole, and Golgi membranes. Our work reveals a conserved helix that governs the localization of VRCs among a group of viruses and points to a possible target for developing broad-spectrum antiviral strategies.

## Introduction

Positive-strand RNA viruses [(+)RNA viruses] are the largest viral class, including many important human and animal pathogens as well as the great majority of plant viruses. Just a few examples include SARS-COV-2 that has infected more than 200 million and led to the death of >4.6 million people in the world (Worldometer.info, 2021); foot and mouth disease virus, the most economically important animal virus; and cucumber mosaic virus (CMV), which infects more than 1,200 plant species. Despite very different host ranges, (+)RNA viruses are grouped into alphavirus-, flavivirus-, and picornavirus-like superfamilies, based on their genome organization and homology of RNA-dependent RNA polymerase (Koonin *et al*, 2015). Each superfamily includes viruses infecting humans, animals, and plants.

A universal feature of (+)RNA viruses is that they assemble their viral replication complexes (VRCs) or replication organelles on specific cellular organelle membranes (Wang, 2015; Paul & Bartenschlager, 2013) and this is attributed to the membrane lipid compositions (Chukkapalli *et al*, 2012; Belov & Kuppeveld, 2012; Zhang *et al*, 2019), among others. The viral replication proteins that commence membrane associations can either be a true integral membrane protein with transmembrane domain(s) (TMDs), such as protein A of Flock House virus (FHV) (Miller & Ahlquist, 2002; Miller *et al*, 2001) or peripheral membrane-associated proteins such as 2C of poliovirus (Teterina *et al*, 2006) and nSP1 of Semliki Forest virus (Lampio *et al*, 2000). Targeting viral replication proteins to the designated organelle membranes is the initial step towards viral replication, and therefore, the knowledge of structural features responsible for protein targeting will not only gain a better understanding of viral replication mechanisms but also underlie our ability to develop new ways for viral control.

Brome mosaic virus (BMV) shares replication features with (+)RNA viruses from many other families and therefore has been useful to investigate fundamental aspects of viral replication and associated processes (Schwartz *et al*, 2002; Diaz & Wang, 2014). BMV is the type member of the family *Bromoviridae* and a member of the alphavirus-like superfamily (Diaz & Wang, 2014; He *et al*, 2021). BMV has a tripartite genome where genomic RNA1 encodes replication protein 1a (1a herein) and RNA2 encodes 2a (2a^pol^ herein) that has a conserved RNA-dependent RNA polymerase domain. RNA3 encodes the movement protein, 3a, and coat protein, which is expressed from RNA4, a subgenomic mRNA that is a replication product of RNA3 (He *et al*, 2021; Wang & Ahlquist, 2008). BMV assembles its spherular VRCs at perinuclear ER (nER) membranes (Schwartz *et al*, 2002). Independent of other viral components, 1a localizes to the nER membranes where it induces the invaginations of the outer nER member into the lumen to form spherules (Schwartz *et al*, 2002), which become VRCs when 2a^pol^ and viral genomic RNA templates are recruited by BMV 1a (Schwartz *et al*, 2002).

BMV 1a has a capping domain at the N-terminus, a helicase-like domain at the C-terminus with a demonstrated NTPase activity (Wang *et al*, 2005), and a linker region enriched in prolines between two domains (He *et al*, 2021). It has been previously revealed that a 113-amino acid region in 1a, region E [amino acids (aa) 367-480], is necessary for its nER membrane association (den Boon *et al*, 2001). Further studies have identified an amphipathic α-helix, helix A (aa 392-409, Fig. 1A) within region E, that regulates 1a’s membrane association, as well as the size and frequency of VRCs, among others (Liu *et al*, 2009). Although helix A was shown to be sufficient to target GFP to membrane fractions, it remained unclear whether helix A is sufficient to target proteins specifically to the nER membrane (Liu *et al*, 2009). A second amphipathic α-helix, helix B (aa 416-433, Fig. 1B), which is within region E and immediately downstream of helix A (Ahola & Karlin, 2015), has been shown to play a crucial role in membrane permeabilization to release the oxidizing potential of the ER lumen, and thus, maintain efficient BMV genome replication (Nishikiori & Ahlquist, 2018).

**Fig. 1.**
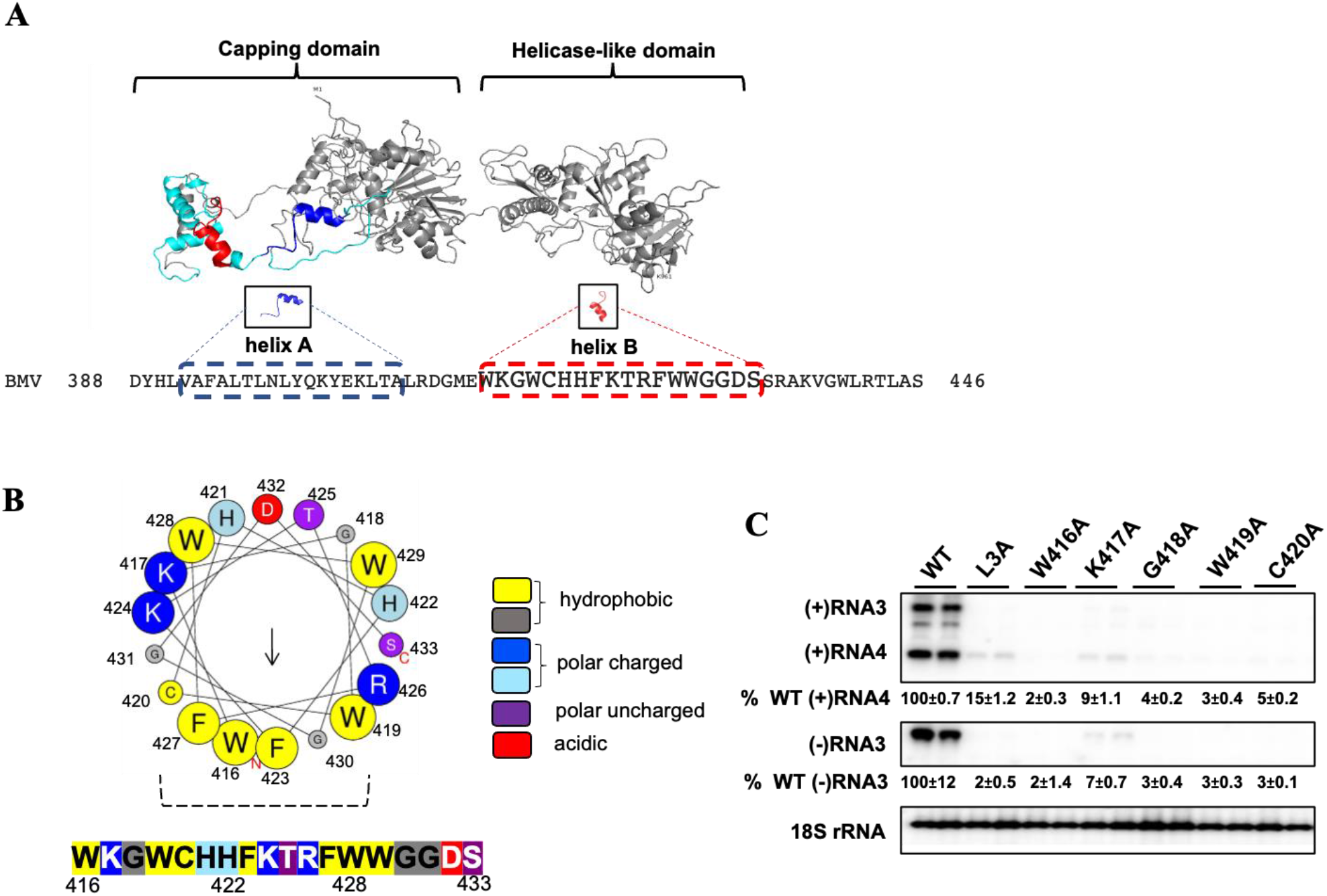
Mutations in BMV 1a helix B inhibit viral genome replication. **(A)** The predicted model for BMV 1a generated using the resolved crystal structure of tomato mosaic virus helicase-like domain as template (PDB-3VKW). Helix A and helix B are highlighted in blue and red respectively and their amino acid sequences are shown. Note region E highlighted in cyan includes helix A and B. **(B)** Helical wheel projection generated using an 18 aa analysis window in Heliquest. The predicted hydrophobic face is formed by W419, G430, F423, W416, F427, and C420, as indicated by the dotted lines. The size of each residue in the helical wheel represents the bulkiness of their sidechain and the arrow in the helical wheel corresponds to the hydrophobic moment. **(C)** BMV genomic replication was measured by using Northern blotting for WT and various 1a mutants. Positive- and negative-strand viral RNAs were detected using BMV strand-specific probes. 18S rRNA served as a loading control. Band intensities were measured using Adobe Photoshop. Note: L3A has 3 Leu residues replaced by Ala in helix A and serves as a negative control.

We report here that helix B is necessary for 1a’s nER membrane association and is sufficient to target several soluble proteins to the nER membrane, extending our understanding on the roles of helix B in targeting 1a to the designated organelle for the VRC formation. Furthermore, we show that the predicted helix B motifs from replication proteins of cowpea chlorotic mottle virus (CCMV), CMV, hepatitis E virus (HEV), and Rubella virus (RuV) direct soluble fluorescent protein(s) to specific organelles where these viruses replicate, revealing a functionally conserved motif across the alphavirus-like superfamily and pointing out a potential antiviral strategy to control a large group of viruses.

## Results

### Helix B of 1a is necessary for BMV RNA replication

For many membrane-associated proteins without TMDs, shallow insertion of amphipathic helices into membranes promotes membrane binding and remodeling (Martyna *et al*, 2017; Drin & Antonny, 2010). BMV 1a does not have any TMD but is tightly associated with the membrane (den Boon *et al*, 2001) at the nER. It has been previously reported that helix A was able to target GFP to membranes based on a membrane flotation assay (Liu *et al*, 2009), however, it was unclear whether helix A is sufficient for 1a’s association with the nER membrane. We tested whether helix B is involved in targeting 1a to the nER membrane or whether a concerted action by both helices is required.

We first tested whether helix B is necessary for viral replication by deleting the entire sequence of helix B to make the 1a-ΔB mutant. In cells expressing 2a^pol^, RNA3, and wild-type (WT) 1a, strong signals were detected for both (−)RNA3 and (+)RNA4, which can only be produced after a full round of replication (Fig. 1C). In parallel with the previous report that deleting helix A blocked BMV replication (Liu *et al*, 2009), no (−)RNA3 or (+)RNA4 band was detected with the 1a-ΔB mutant, indicating that helix B is necessary for BMV replication (Table 1).

**Table 1.**
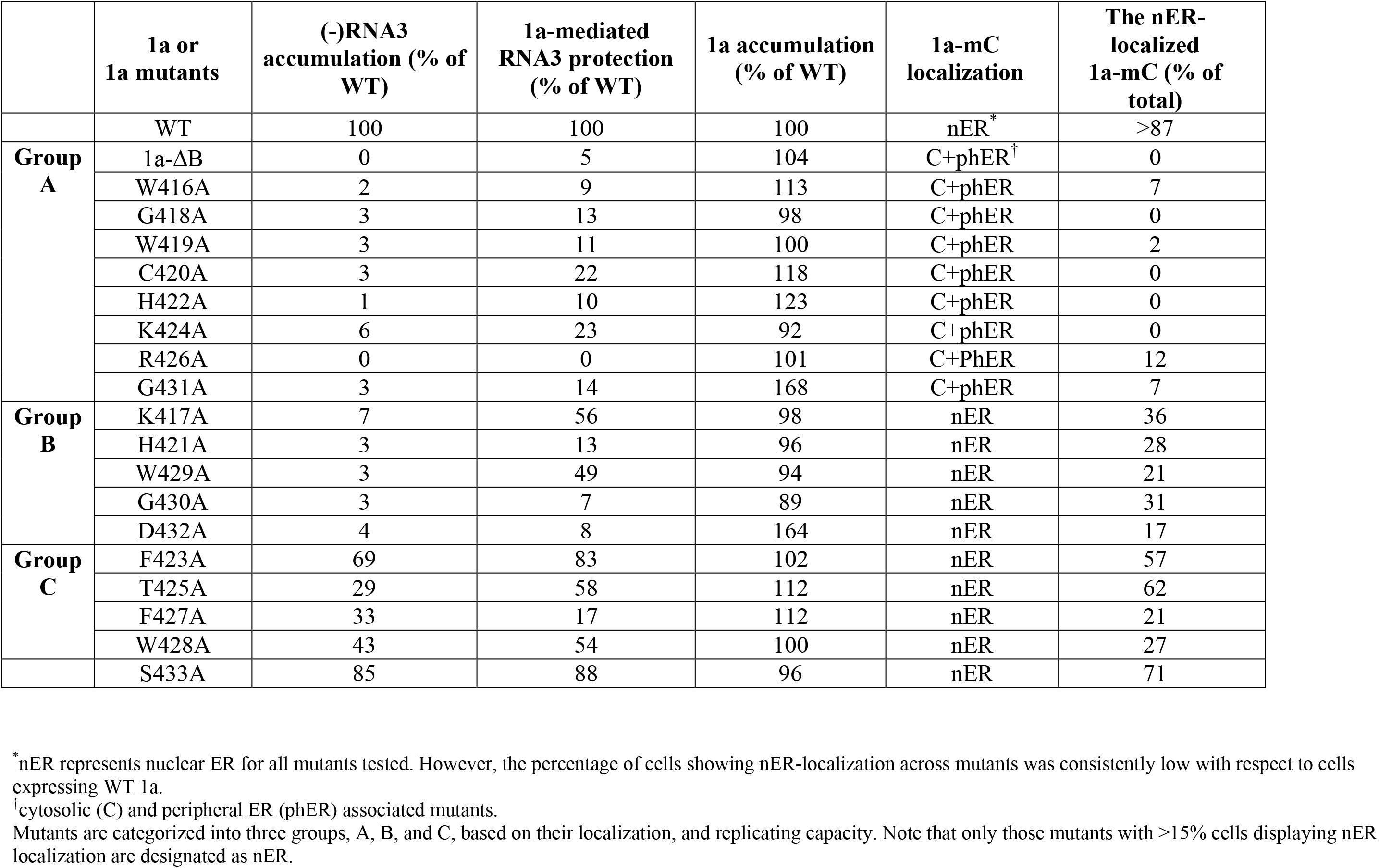
Characteristics of 1a helix B mutants.

We next substituted every amino acid in helix B with alanine and characterized each mutant on their ability to support BMV RNA replication. Most mutants failed to support genome replication, producing (−)RNA3 at levels less than 10% of that with WT 1a (Table 1 and Fig. 1C). These mutants were comparable to that of a helix A mutant, L3A, which has 3 leucine residues replaced by alanine, is not associated with the nER membrane, and is defective in supporting the viral replication (Fig. 1C) (Liu *et al*, 2009). Only 5 mutants in Group C (F423A, T425A, F427A, W428A, and S433A) produced (-)RNA3 that were higher than 25% of WT levels. The inhibited BMV RNA replication was not due to the affected expression and/or stability by each mutation, because each mutant accumulated at similar levels to WT 1a (Table 1). These data indicated that helix B is necessary for viral replication processes.

### Helix B is necessary for the nER membrane association of BMV 1a

To assess the impact of each mutation on 1a’s membrane association, we performed a membrane flotation assay by an iodixanol density gradient. Membrane flotation assays distinguish membrane-associated or -integral proteins from those that are soluble or cytoplasmic. Using the established conditions where membranes and membrane-proteins float atop the gradient, and soluble fractions sink to the bottom (den Boon *et al*, 2001), we confirmed that WT 1a and membrane-protein dolichol-phosphate mannosyltransferase (Dpm1) were both detected primarily in the top two fractions, and the soluble protein phosphoglycerate kinase (Pgk1) was in the bottom two fractions (Fig. 2A). Membrane association was determined as the percentage of a protein in the top three fractions against the sum of all 6 fractions (Fig. 2A). About 88% of WT 1a, 96% of Dpm1, and 6% of Pgk1 were detected in the top 3 fractions (Fig. 2A).

**Fig. 2.**
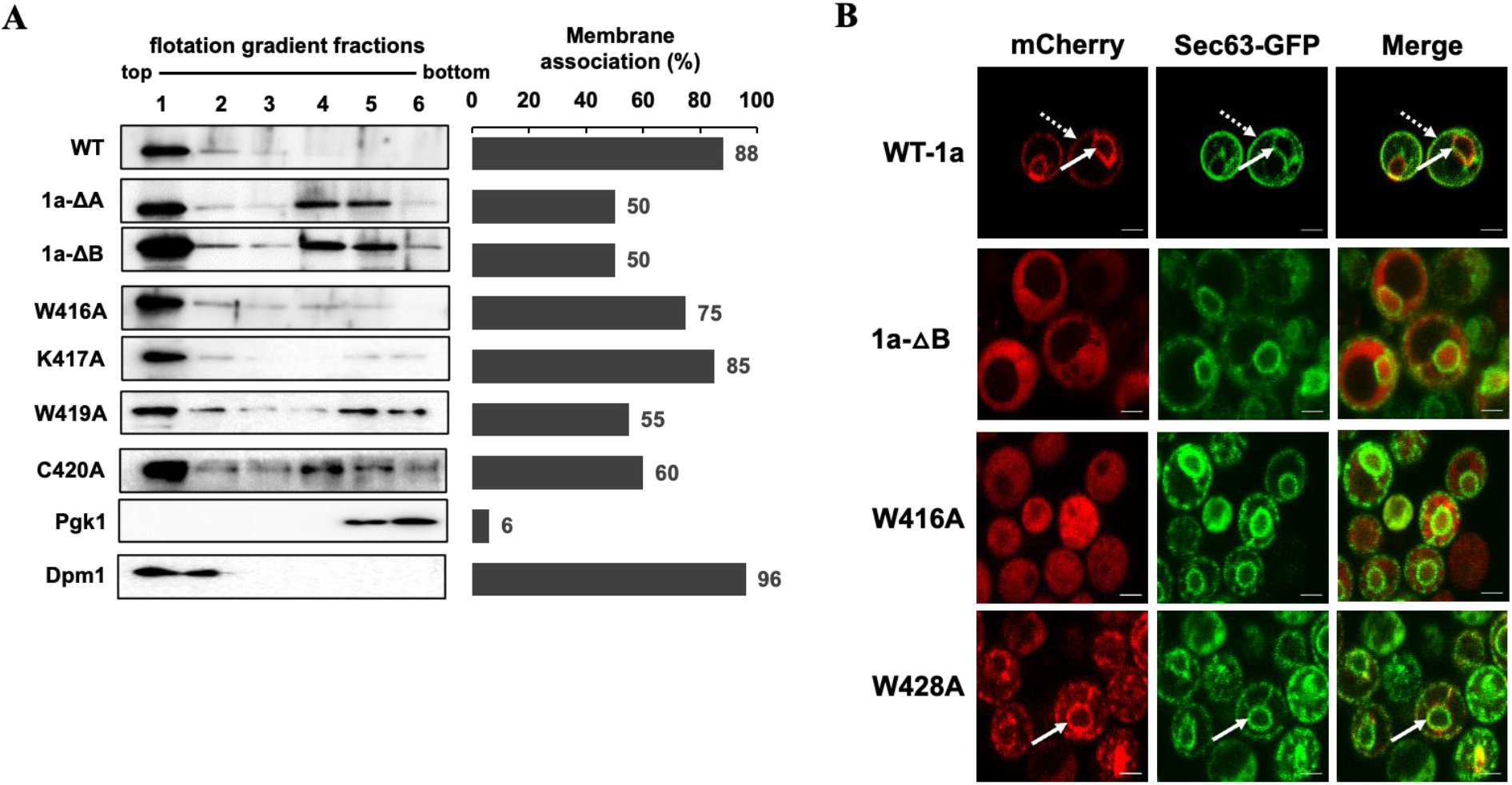
Mutations in BMV 1a helix B inhibit the nuclear ER membrane association of BMV 1a. **(A)** Membrane association of BMV 1a mutants was determined via a membrane flotation assay. Histograms represent the percentage of membrane-associated proteins as determined by the signal intensity of each protein segregated into the top three membrane fractions compared to the sum of all six fractions. Lysates of yeast cells expressing WT or mutant 1a were subjected to an iodixanol density gradient and analyzed by Western blotting using antibodies against ER membrane protein Dpm1, soluble protein Pgk1, or His6 for His6-tagged BMV 1a derivatives. **(B)** Localization of mCherry-tagged 1a mutants in yeast cells co-expressing Sec63-GFP as examined using confocal microscopy. Nuclear ER and peripheral ER associations are indicated by solid and dotted arrows, respectively. The yellow signal in the merge panel is representative of colocalization of mCherry-tagged WT or mutant 1a with Sec63-GFP. Scale bars: 2μm.

We chose five replication defective mutants for the membrane flotational analysis, including 1a-ΔB with helix B deleted, W416A, W419A, C420A, and K417A. The mutant W416A, W419A, C420A form the predicted hydrophobic face of the amphipathic helix (Fig. 1B) and also present at the beginning of the helix sequence. The mutant K417A was chosen due to its charged nature and potential contribution to membrane association. The 1a-ΔB mutant showed a 50% accumulation in the membrane fractions. The other 50% accumulated in the cytosolic fractions, with most accumulated in fractions 4 and 5, similar to the previously characterized deletion mutant, 1a-ΔA (Liu et al, 2009). Mutants W416A and K417A showed a strong membrane association with 75% and 85% of total protein in the membrane fractions, whereas W419A and C420A showed membrane association of about 55% and 60%, respectively (Fig. 2A).

Since membrane flotation results are not informative in terms of specific organelle membranes where target proteins reside, we checked the localization for all 1a mutants using fluorescence and/or confocal microscopy. Since 1a predominantly localizes to the nER membrane when expressed alone (Restrepo-Hartwig & Ahlquist, 1999), we expressed 1a helix B mutants with a C-terminally fused mCherry (mC) in yeast cells along with an ER protein marker, Sec63, with a C-terminally fused GFP. Since proteins could obtain distinct conformation due to the presence of an additional protein moiety (Argos, 1990; Chen *et al*, 2013), we incorporated a 32 amino acid linker (herein L32) (Finnigan *et al*, 2016) with desirable small polar amino acids, such as Thr, Ser, and Gly, between the carboxy-terminal of the protein and mC. The mC-tagged WT 1a protein was efficiently localized to the nER (the inner ring pointed by solid arrows in Fig. 2B) and the peripheral ER regions (the outer ring as pointed by dotted arrows). When helix B was deleted from 1a, the mC signal was throughout the cytoplasm, indicating that helix B was necessary for 1a’s association with the nER membrane (Fig. 2B). Helix B mutants with single substitutions that were defective in replication were segregated into two sets based on their ability to localize to the nER membrane, as shown for a representative mutant for each set in Fig. 2B. Mutants in Group A (W416A, G418A, W419A, C420A, H422A, K424A, R426A, and G431) were detected mostly throughout the cytosol (Fig. 2B, Table 1). Group B comprised of mutants that could partially localize to the nER membrane, including K417A, H421A, F423A, T425A, F427A, W428A, W429A, G430A, D432A, and S433A. Although Group B mutants retained their ability to associate with the nER membrane, there were not as many cells showing nER localization as observed in cells expressing WT 1a protein (Table 1). We also noticed puncta in mutants F423A, T425A, and W429A (Supplemental Fig. EV2 C).

### Mutations in helix B inhibit 1a-mediated RNA3 protection

BMV 1a, in the absence of other viral components, induces the formation of spherules in yeast cells. 1a also recognizes and recruits viral genomic RNA and 2a^pol^ into spherules, which become VRCs (Schwartz *et al*, 2002). When expressed alone, RNA3 is unstable in yeast cells with a half-life of 3-10 minutes, whereas in the presence of 1a, the half-life of RNA3 increases to more than 3 hours, and the accumulated RNA3 increases 5-20 folds, depending on the levels of expressed 1a (Janda & Ahlquist, 1998; Wang *et al*, 2005). As shown in Fig. EV3, an 11-fold increase of RNA3 was observed in the presence of WT 1a but not in the 1a-ΔB mutant, indicating the importance of helix B in 1a-mediated RNA3 protection. In addition, the majority of helix B mutants with a single substitution showed a comparatively lower amount of protected RNA3 than WT 1a (Fig. EV3, Table1). As shown in Fig. EV3 for W416A, W418A, W419A, and C420A, all mutants in Group A showed <25% levels of the accumulated RNA3 in cells expressing WT 1a (Table 1), consistent with the fact that all mutants in Group A failed to associate with the nER (Table 1), and thus, these mutations may have substantially interfered with 1a’s ability to form spherules. However, this note should be confirmed in the future with electron microscopy. The accumulated RNA was widely varied among mutants in Group B and C, which were partially associated with the nER membrane, from 17% to 88% of WT levels (Table 1), as represented by K417A at 56% of WT 1a levels (Fig. EV3).

### Helix B is sufficient to target soluble proteins to the nER membrane

It was previously reported that helix A alone or the entire region E is capable of targeting GFP to the membrane fractions based on membrane flotation assays (Liu *et al*, 2009). To estimate the membrane association capacity of helix B in comparison to the entire region E or helix A, we performed a membrane flotation assay using lysates from cells expressing GFP-tagged region E, helix A, helix B, or helices A and B.

Consistent with previous studies, when fused to region E or helix A, 54% and 40% of GFP were detected in the membrane fractions compared to the sum of GFP signal across all fractions, respectively (Liu *et al*, 2009; den Boon *et al*, 2001). Cells expressing both helices showed 46% of GFP in the membrane fractions but GFP was detected across all the collected fractions, similar to cells expressing region E-GFP. Surprisingly, 68% of helix B-GFP was associated with the membrane fractions, suggesting that helix B is critical in targeting 1a to membranes (Fig. 3A).

**Fig. 3.**
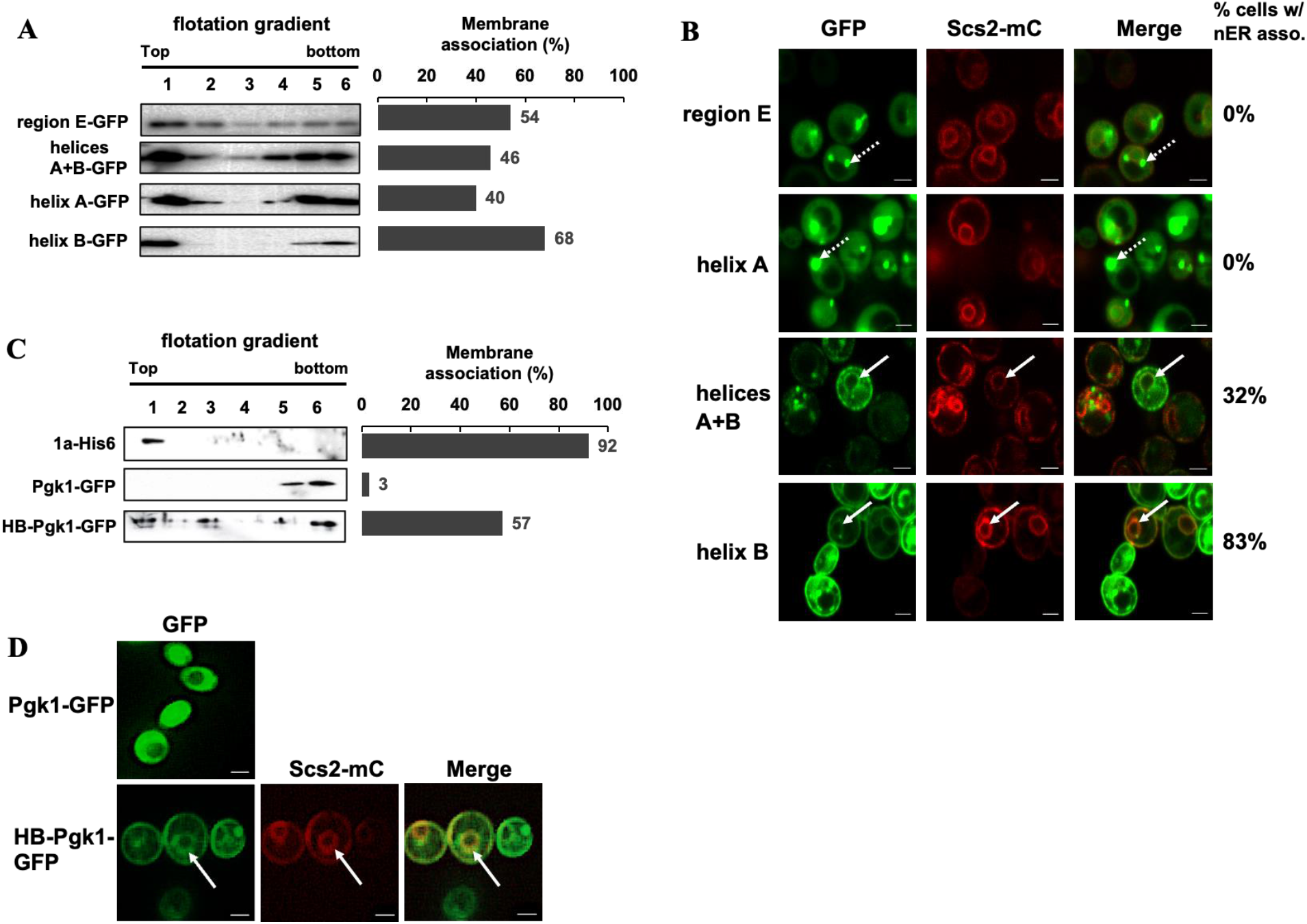
Helix B is capable of targeting soluble proteins to the nuclear ER membrane. **(A)** Membrane flotation analysis of GFP-fused region E, helices A+B, helix A, and helix B. Histograms represent the percentage of proteins detected in the top three fractions of the flotation gradient. A representative western blot is shown, with bands corresponding to respective 1a fragments detected using anti-GFP antibody. **(B)** Localization of GFP-tagged region E, helices A+B, helix A, or helix B co-expressed with a mCherry-tagged ER marker protein Scs2. Dotted arrows point to punctate structures and solid arrows point to the nER membrane. Images were obtained using a confocal microscope. **(C)** Membrane flotation analysis for Pgk1-GFP and HB-Pgk1-GFP, using 1a-His6 as a positive control. **(D)** Localization of HB-PGK-GFP at the nER membranes with an ER marker Scs2. Solid arrows point to the nER membrane. Images are obtained using a fluorescence microscope. Scale bars: 2μm.

Since membrane fractions represent many cellular organelle membranes, we turned to microscopy to determine whether helix B-GFP is associated with the nER membrane specifically. To assess whether helix B-GFP is associated with the nER association, we expressed helix B-GFP along with a mCherry-tagged ER marker Scs2. We also included helix A and region E for comparative analysis. Each was expressed under the control of the *GAL1* promoter, and the cells were harvested across various time points post induction when the carbon source was switched from raffinose to galactose. Region E did not localize GFP to the nER membrane across various time points but aggregated as puncta in cells, consistent with the previous report (den Boon *et al*, 2001). These punctate structures that were observed at 6-7 hours post-induction became more diffused into the cytosolic region at 12h post-induction. Helix A-GFP primarily showed a rather peripheral and dispersed localization along with strong puncta (Fig. 3B). Cells expressing a 1a fragment encompassing both helices A and B showed a partial nER association in ~32% of the cells. Helix B-GFP, agreeing well with the membrane flotation results (Fig. 3A), was detected as a two-ring pattern typical of nER and peripheral ER association in nearly 83% of the cells (Fig. 3B). On the contrary, none of the cells expressing either region E-GFP or helix A-GFP showed the fluorescent signal in the nER membrane (Fig. 3B), suggesting that helix B is not only sufficient for the nER targeting but also represents the strongest targeting information in region E. Shorter helix B sequences spanning only the first seven (W416-H422) or eleven (W416-R426) amino acids of the alpha-helix B failed to associate with the nER (Fig. EV2 B), suggesting that the amphipathicity is governed by more than the first eleven residues of helix B.

To further determine the capacity of helix B in targeting other soluble proteins to the nER membrane, we fused helix B at the N-terminus of an endogenous yeast cytosolic protein. We chose yeast Pgk1, a soluble protein that is primarily detected in the cytoplasm when fused with GFP (Fig. 3D) and readily detected at the bottom-most fractions in our flotational analysis in the absence (Fig. 2A) or presence of GFP (Fig. 3C). The fusion protein, HB-Pgk1-GFP gained the ability to associate with membranes and 57% of protein was detected at the top three fractions (Fig. 3C). In agreement with the membrane flotation results, it could also be seen localized to the nER and peripheral ER regions, colocalizing with Scs2-mC (Fig. 3D), similar to WT-1a protein that brought mCherry to the nER membrane (Fig. 2B). Collectively, our results indicated that Helix B acted as a dominant targeting peptide that directed soluble proteins to the nER membrane.

### The amphipathic α-helix represented by helix B is functionally conserved among viruses in the alphavirus-like superfamily

BMV is a representative member of the alphavirus-like superfamily. Viruses in this superfamily contain a methyltransferase-guanylyltransferase (MTase-GTase) domain whose activity has been experimentally confirmed in the nsP1 of SFV (Laakkonen *et al*, 1994; Scheidel *et al*, 1989), 1a of BMV (Kong *et al*, 1999), and open reading frame 1 (ORF1) of hepatitis E virus (HEV) (Magden *et al*, 2001). Among the alphavirus-like superfamily, a conserved “Iceberg” region has been identified at the C-terminus of the viral MTase-GTase domains with the presence of two amphipathic helices (Ahola & Karlin, 2015), similar to the helices A and B of 1a.

Since amphipathic helix B is a predicted secondary structure conserved across the alphavirus-like superfamily (Ahola & Karlin, 2015) with the actual predicted boundary of each helix varying across the superfamily, we wanted to test whether the predicted helix B from other members is also functionally conserved in terms of targeting the replication protein to specific organelle membranes. We noticed that the primary sequence conservation is retained for members in the same genus, such as BMV and CCMV (Sibert *et al*, 2018), whereas the conservation is not high across members of a family, including BMV and CMV, where BMV is a type member of the genus Bromovirus and CMV is of the genus Cucumovirus (Supplemental Fig. EV1). The primary sequence conservation of helix B was even lower for human viruses such as HEV and RuV when compared against BMV helix B (Ahola & Karlin, 2015).

We fused the predicted helix B of CCMV, CMV, HEV, and RuV to the N-terminus of fluorescent proteins and tested the localization of each recombinant protein in yeast cells, along with an ER marker, Scs2-mC or Sec63-GFP. The CCMV helix B, when fused to GFP (CCMV-HB-GFP), was capable of targeting GFP to the nER membrane and colocalized with Scs2-mC (Fig. 4A). This agreed well with the previous report that CCMV replicates in the nER membranes (Sibert *et al*, 2018; Kim, 1977) and CCMV 1a, when expressed in yeast cells in the absence of other viral proteins, induces the formation of spherular invaginations at the outer nER membranes (Sibert *et al*, 2018).

**Fig. 4.**
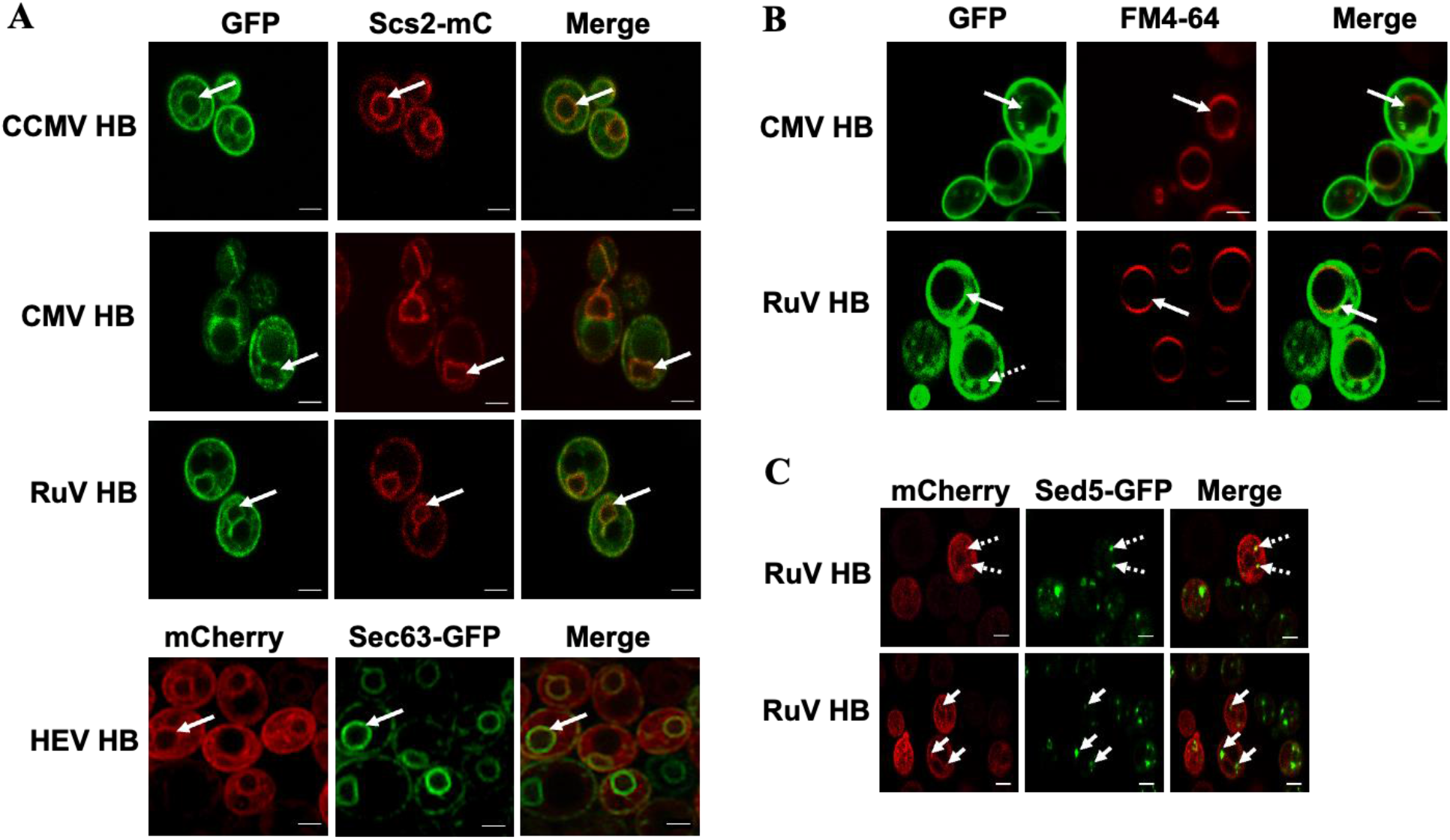
Helix B-mediated organelle targeting is conserved across members of the alphavirus-like superfamily. **(A)** Colocalization of GFP-tagged HB of CCMV, CMV, or RuV with an ER marker Scs2-mC, and colocalization of the mCherry-tagged HEV HB and Sec63-GFP at the nER membrane. Solid arrows point to the nER membrane. Note arrows in second and third rows (CMV and RuV) distinguish the nER membrane from other membranous regions present in the cells. **(B)** Colocalization observed for CMV HB and RuV HB with vacuolar membranes stained by a lipophilic dye, FM4-64. The solid arrows in both panels point to vacuolar membranes. The dotted arrow in the RuV-HB panel shows punctate structures. **(C)** Colocalization of RuV HB-mC with Sed5-GFP, a marker for vesicles that traffic from the ER to the Golgi. Images were obtained using a confocal microscope. Scale bars: 2μm.

Unlike BMV and CCMV, CMV replicates in the vacuolar membrane of plant cells, the tonoplast (Hatta & Francki, 1981; Wang *et al*, 2021b). Interestingly, CMV helix B targeted GFP to the nER and vacuolar membranes in yeast cells (Fig. 4A). At 6-7 hours post induction of its expression, CMV-HB-GFP localized at the nER in 64% of cells and at vacuoles in 77% of cells, 12% of cells showed both nER as well as vacuolar localization (n=175). Vacuolar localization of CMV-HB-GFP dominated when grown beyond 7 hours in induction media. It is possible that CMV HB-GFP was localized first to ER membranes, following the protein secretory pathway, before reaching vacuolar membranes.

HEV is a non-enveloped (+)RNA virus in the family of *Hepeviridae* (Kenney & Meng, 2019). It is one of the leading viral pathogens causing acute hepatitis. HEV encodes 3 major ORFs. ORF1 is a polypeptide that is processed by viral encoded proteases. Two amphipathic alpha-helices, helix A (aa 362-380) and B (aa 399-417), are also present in the Y domain (aa 216-442) of ORF1. When fused to GFP, Helix B of HEV ORF1 also colocalized with the ER marker Sec63 at the nER membrane (Fig. 4A), which is the purported site of HEV replication (Kenney & Meng, 2019). However, mCherry signal was also found between the nER and peripheral ER, similar to tubular ERs that connect the nER and peripheral ER.

RuV is known to utilize multiple organelle membrane sites for its replication. These membranes include those of Golgi, endosomes, vesicles, and vacuoles of endolysosomal origin in addition to ER (Fontana *et al*, 2007). The viral NS protein P150 is sufficient for organelle targeting (Kujala *et al*, 1999) and also contains a Y domain similar to HEV with the presence of two amphipathic helices. We expressed RuV helix B with a C-terminally fused GFP and confirmed its localization at the vacuolar regions using a lipophilic dye, FM4-64 that stains the vacuolar membrane (Fig. 4B), and its localization at the nER by using the ER marker Scs2-mC (Fig. 4A). We also fused RuV HB to mC and coexpressed RuV-HB-mC along with an early Golgi compartment marker Sed5 that was fused with GFP. As shown in Fig. 4C, RuV-HB-mC did colocalize with the punctate structures seen throughout the cytoplasm labeled by Sed5-GFP.

### Helix B from BMV and CMV target fluorescent proteins to ER membranes and tonoplast in plant cells

The ability of BMV and CMV helix B to target GFP to the nER and/or vacuole membranes in yeast cells prompted us to check whether these helices could do the same *in planta*. To do this, we fused BMV or CMV helix B to the N-terminus of YFP in a T-DNA binary vector under the control of the cauliflower mosaic virus (CaMV) 35S promoter (Zhao *et al*, 2020) and expressed by agroinfiltration in histone 2B-RFP (H2B-RFP) transgenic *Nicotiana benthamiana* plants (Martin *et al*, 2009). The localization of YFP signal in *N. benthamiana* cells was observed under a laser confocal microscope at 40 hours post agroinfiltration (hpai), which is the commonly used time point to observe transiently expressed proteins (Zhao *et al*, 2020). While free YFP was observed in the cytoplasm and the nucleus, BMV HB-YFP signal was observed at the nER membranes, surrounding the histone 2B-RFP-represented nucleus (Fig. 5A). To confirm BMV HB-YFP was associated with the nER membrane, we co-expressed BMV HB-YFP and an mC-tagged ER marker (Nelson *et al*, 2007). The YFP signal was co-localized with the mC signal in the nER and peripheral ER (Fig. 5B). In addition, we have observed multiple ER-localized aggregates that were not present in cells expressing YFP (Fig. 5A). Surprisingly, we also observed small vesicle-like structures that contain both signals of YFP and mC (arrowheads in Fig. 5B), indicating the ER origin of these vesicles. However, the possible functions of these vesicle-like structures are unclear.

**Fig. 5.**
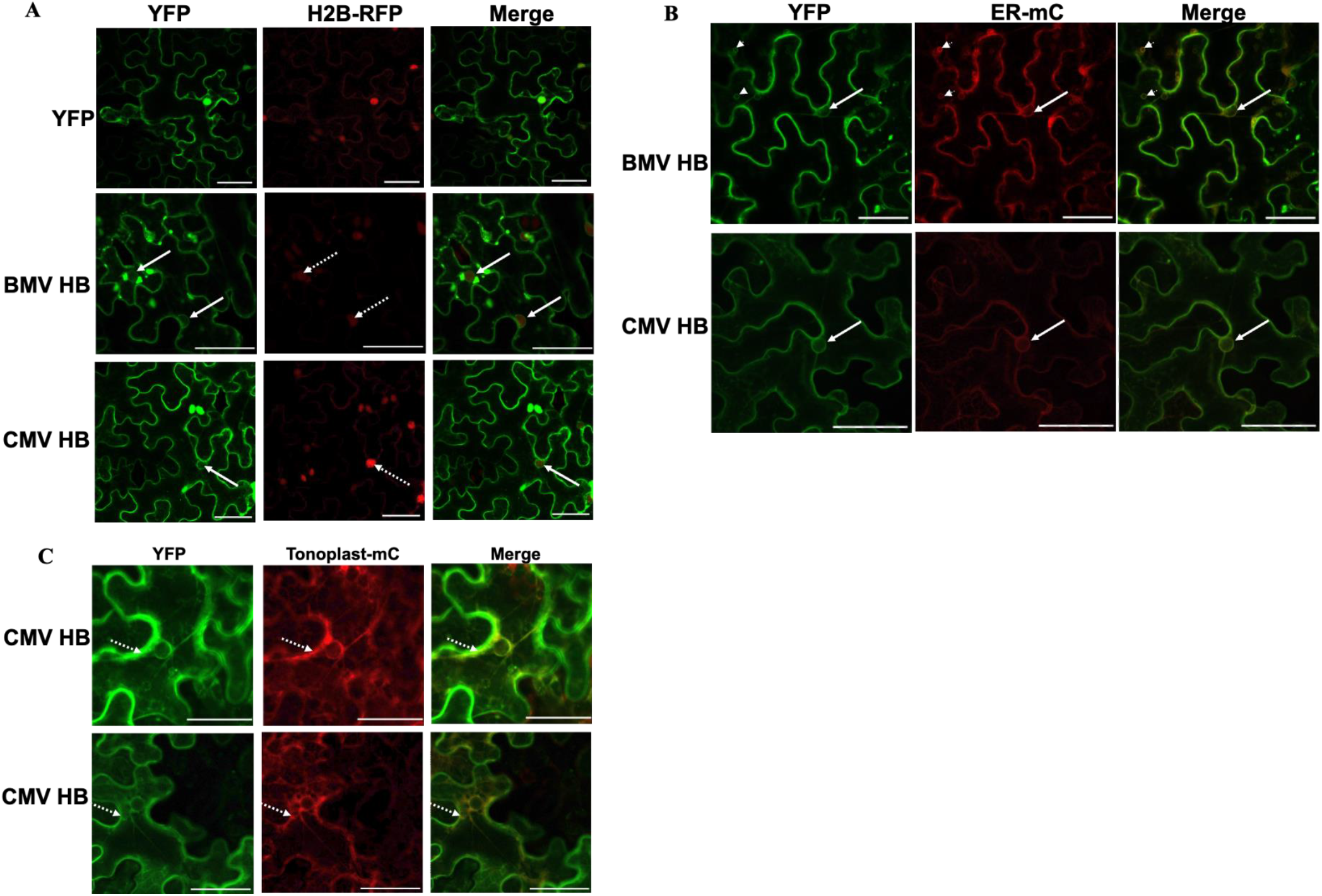
Localizations of BMV and CMV helix B in *N. benthamiana* cells parallel those in yeast cells. **(A)** YFP-tagged Helix B from BMV or CMV was expressed in histone 2B-RFP transgenic *Nicotiana benthamiana* plants via agroinfiltration. Solid and dotted arrows point to the nER membrane and the nucleus, respectively. **(B)** BMV HB-YFP was co-expressed with a mCherry-tagged ER marker. Arrows point to the nER membrane and arrowheads point to small vesicles that are colocalized with the ER marker. **(C)** Colocalization of CMV HB-YFP with a mCherry-tagged tonoplast marker as indicated by dotted arrows. The fluorescence signal was observed under laser confocal microscopy at 40 hpai. Scale bars: 50μm.

We next tested the localization of YFP-tagged CMV HB in H2B-RFP-transgenic *N. benthamiana* cells. Similar to what we have observed in yeast cells, CMV HB-YFP was localized at the nER membrane surrounding the nucleus and co-localized with the ER marker (Fig. 5A and B). Agreeing well with the report that CMV 1a, when expressed alone, localizes to vacuolar membranes (Wang *et al*, 2021b), CMV HB-RFP was detected primarily at the tonoplasts as it colocalized with an mC-tagged tonoplast marker (Fig. 5C). These results confirmed the results observed in yeast that CMV HB can target YFP to both ER and vacuolar membranes. We also included YFP-tagged CCMV HB in our study and found nER localization in these cells (Supplemental Fig. EV4), agreeing well with the results in yeast cells (Fig. 4A).

## Discussion

The membrane association of viral replication complexes is a key feature across all (+)RNA viruses. Different (+)RNA viruses replicate in specific organelle membranes. BMV, CCMV, and beet black scotch virus replicate on ER membranes, whereas CMV, tomato bushy stunt virus, barley stripe mosaic virus, and turnip crinkle virus prefer membranes of the tonoplast, peroxisome, chloroplast, and mitochondria, respectively (Kim, 1977; Hatta & Francki, 1981; Cao *et al*, 2015; McCartney *et al*, 2005; Wang *et al*, 2021a; Blake *et al*, 2007; Restrepo-Hartwig & Ahlquist, 1996). Some plausible reasons for the varied preference of membranes are believed to be due to the membrane lipids and/or protein compositions (Zhang *et al*, 2012; Lee *et al*, 2001; Lee & Ahlquist, 2003; Zhanga *et al*, 2016; Zhang *et al*, 2018). However, it is still unclear what structural determinant(s) within the viral replication proteins are responsible for targeting these proteins to specific organelles, especially for those that lack TMDs, such as BMV 1a. We report here that a single amphipathic α-helix in 1a, helix B, plays a key role in determining 1a’s association with the nER membrane. Helix B is necessary for 1a’s nER membrane association and is sufficient to target several soluble proteins to the nER membrane, extending our understanding beyond its role as to regulates the oxidizing potential of the ER lumen (Nishikiori & Ahlquist, 2018). We further show that a similar helix in the replication proteins of CCMV, CMV, HEV, and RuV is able to target fluorescent proteins to the specific organelles where their VRCs are formed, revealing a conserved feature among members of the alphavirus-like superfamily.

### BMV 1a helix B is necessary and sufficient for 1a’s association with the nuclear ER membrane

Many viral replication proteins from (+)RNA viruses contain TMDs that target them to the designated organelle membranes for initiating the VRC formation. Protein A of FHV, when expressed alone, is targeted to the outer mitochondrial membrane where FHV forms VRCs (Miller & Ahlquist, 2002; Ertel *et al*, 2017). Protein A has an N-terminal TMD (aa 1-46) that is sufficient to target GFP to mitochondrial membranes (Miller & Ahlquist, 2002). Replication proteins of some (+)RNA viruses lack a TMD but possess amphipathic α-helix or helices, includes BMV 1a (helix A, aa 392-407 and helix B aa 416-433) (Liu *et al*, 2009; Nishikiori & Ahlquist, 2018; Ahola & Karlin, 2015), Chikungunya virus nsP1 (aa 244-263) (Gottipati *et al*, 2020), poliovirus 2C (aa 19-36) (Teterina *et al*, 2006; Laufman *et al*, 2019), Hepatitis C virus 5A (aa 7-28) (Elazar *et al*, 2003), and SFV nsP1 (aa 245-264) (Spuul *et al*, 2007), among others.

Amphipathic α-helices have long been recognized as generators and/or sensors of membrane curvature (Shen *et al*, 2012; Drin & Antonny, 2010). Residues on the hydrophobic face of the helix insert into the fatty acid tails while polar residues are involved in electrostatic interactions with the polar head of phospholipids (Jensen *et al*, 2011). It has long been understood that the amphipathic α-helix serves as an anchor, inserting TMD-lacking viral proteins shallowly into the membranes. Our recent data indicated that the helix of poliovirus 2C played an active role in remodeling membranes. In an *in vitro* tubulation assay, the synthesized peptide representing the helix of poliovirus 2C converted large spherical liposomes into tubules and smaller vesicles that were stable for a week (Varkey *et al*, 2020). In addition, mutations disrupting the helicity of the 2C helix inhibited membrane remodeling *in vitro* when expressed as a peptide and blocked viral replication when incorporated into full-length viral genome (Varkey *et al*, 2020).

Helix B of BMV 1a was first predicted by Ahola and Karlin while analyzing BMV 1a, SFV nSP1, and equivalent replication proteins among the alphavirus-like superfamily (Ahola & Karlin, 2015). The predicted helix B was further characterized for its amphipathicity and its viroporin activity that allows oxidizing environment within BMV VRCs (Nishikiori & Ahlquist, 2018). In this study, we determined that helix B is necessary and sufficient for BMV 1a’s association with nER membrane, and thus, essential for BMV genomic replication (Table 1). The vast majority of the helix B mutants (group A and B) with an Ala substitution either failed or supported no more than 10% of WT 1a genomic replication (Table 1), indicating a crucial role for helix B in this process. This also aligns with the reported helix B mutants K424G and R426K for selectively lacking viroporin activity that failed to replicate (Nishikiori & Ahlquist, 2018).

The 19 helix B mutants generated in this study can be divided into three groups based on genomic replication and their localization. Group A contains nine mutants, including the helix B deletion mutant, and eight Ala-replacement mutants (W416A, G418A, W419A, C420A, H422A, K424A, R426A, and G431A), which were primarily detected in the cytoplasm and possibly peripheral ER membranes (Table 1). Future experiments will further address to which organelle(s), other than the nER, these 1a mutants were targeted. Group B includes five mutants that were associated with the nER at lower efficiency than WT, ranging from 17% to 36% of cells (Table 1). For instance, K417A was detected in the nER membrane in 36% of cells, compared to 87% of cells with WT 1a (Table 1). Mutants in both Groups A and B supported less than 10% of viral genomic replication compared to WT 1a. Another five mutants supporting partial replication and inefficiently localized at the nER membrane are assigned to group C. It is possible that some mutants were only transiently associated with the nER. Given the fact that these mutants accumulated at a similar level to WT (Table 1), we concluded that helix B is necessary for BMV 1a’s nER membrane association and thus, genome replication.

Soluble proteins, GFP (Fig. 3B), mC (Supplemental Fig. EV2 A), and Pgk1 (Fig. 3D), were targeted primarily to the nER and also peripheral ER membrane when fused to helix B at their N-terminus. However, helix A-tagged GFP did not (Fig. 3B), indicating that BMV 1a helix B is responsible for 1a’s targeting to the nER membrane. We are unable to determine whether helix B-mediated nER targeting is due to the specific lipid composition of the organelle membranes or whether it interacts with a specific host protein that localizes to the nER (Li *et al*, 2016). It will be exciting to study in the future whether targeting is an active process requiring host machinery in trans or whether it is a passive process relying on cis-elements within the helix and their internal propensity to bind the lipid bilayer.

### Helix B-mediated organelle targeting is a conserved feature across members within the alphavirus-like superfamily

The initial step for a successful viral infection is to assemble functional VRCs at the target organelle. This starts with targeting viral replication proteins to the organelle where VRCs will be established. Disruption of this step will block viral replication at its early stage. For instance, we have reported a mutant of yeast protein Cho2p (choline requiring 2), *Cho2p-aia*, that retargeted BMV 1a away from the nER and ended as peripheral ER-localized puncta, and thus, significantly inhibited BMV replication (He *et al*, 2019).

Knowing that helix B of 1a is sufficient to target soluble proteins to the site of BMV replication, we tested the conservation of this feature across the members of the alphavirus-like superfamily, given the fact that two similar amphipathic α-helices have been identified based on sequence analysis with helix B immediately downstream of helix A (Ahola & Karlin, 2015). Helix B from CCMV, which is in the same genus as BMV and replicates in the nER membrane (Sibert *et al*, 2018), targeted mC to the nER membrane, as hypothesized (Fig. 4A). Helix B from CMV 1a targeted GFP to vacuolar membranes, similar to the tonoplast where CMV replicates in plant cells (Hatta & Francki, 1981; Wang *et al*, 2021b). Importantly, the results for the helices from BMV, CMV, and CCMV were similar in both yeast and plant cells (Figs. 4 and 5). We further demonstrated that predicted amphipathic α-helices in HEV and RuV were able to target mCherry to the sites of replication, the ER membrane for HEV, and Golgi, ER, and vacuolar membranes for RuV (Fontana *et al*, 2007; Kenney & Meng, 2019) (Fig. 4). Our work, therefore, demonstrates that helix B in the replication proteins of a group of diverse viruses in the alphavirus-like superfamily is indeed a conserved determinant for targeting the viral replication proteins to their specific sites of replication.

It has been reported that the amphipathic α-helix (aa 7-28) in HCV 5A was able to target GFP to ER membranes, where HCV assembles its VRCs (Wang & Tai, 2016). For poliovirus 2C, a 38 amino acid-long fragment (amino acid 1-38) encompassing the amphipathic α-helix (amino acid 19-36) was able to associate with lipid droplets, where poliovirus VRCs acquire fatty acids for VRC formation (Laufman *et al*, 2019). With two amphipathic helices (αA and αB) that are separated by several amino acids, the 41 amino acid-long chloroplast targeting domain (CTD) of the 140K/98K replication protein from turnip yellow mosaic virus (TYMV), another virus from the alphavirus-like superfamily, targets GFP to the outer chloroplast membranes (Moriceau *et al*, 2017). These results have pointed out the role of amphipathic α-helix from individual viruses in targeting specific organelles. Our work, going further, systematically analyzed a range of helices from replication proteins of viruses in the alphavirus-like superfamily and confirmed a conserved role of these helices in targeting viral replication proteins to specific organelle membranes where VRCs are formed.

### Different BMV 1a helices may play different roles in the VRC formation

The helix A peptide of 1a becomes helical when incubated with lipid bilayer-mimicking micelles (Liu *et al*, 2009). Mutations in helix A generated two groups of mutants that were all defective in supporting BMV genome replication. Group I mutants did not associate with the nER membrane, indicating a necessary role of helix A in 1a’s nER association. Mutants in group II induced the formation of VRCs that were more abundant but smaller in size compared to those induced by WT 1a, pointing out a possible role in generating or facilitating the remodeling of membranes into VRCs (Liu *et al*, 2009). Helix A was able to target GFP to membrane fractions in the membrane flotation assay (Liu *et al*, 2009), however, our data showed that helix A was not able to target proteins to the nER on its own (Fig. 3B). Further, helix A and B helices together were not as effective as helix B alone in targeting GFP to the nER membrane (Fig. 3B). It is possible that the presence of helix A might affect the conformation of helix B. Based on the data from Liu et al (Liu *et al*, 2009) and our results, we propose that helix A may participate in membrane remodeling but not 1a targeting and helix B is responsible for 1a’s targeting to the nER membrane.

In this regard, the roles of BMV 1a helix A and B are different from αA and αB in the CTD of TYMV 140K/98K replication protein. TYMV is in the Tymo group of the alphavirus-like superfamily, different from the Alto group that includes BMV, CCMV, CMV, HEV, and RuV (Ahola & Karlin, 2015). TYMV replicates at the outer membrane of chloroplasts and its replication protein 140K/98K is responsible for targeting and remodeling the chloroplast membranes (Moriceau *et al*, 2017), similar to the roles of BMV 1a. The CTD is able to target GFP to the chloroplast membranes. However, it has been clearly demonstrated that disrupting the helicity of either αA or αB does not affect the CTD’s ability to target GFP to the chloroplast membrane, indicating either helix is sufficient when present in the CTD. This is in contrast to the BMV 1a helices A+B domain that showed much reduced efficiency in targeting GFP to the membrane fraction (Fig. 3A) and the nER membrane (Fig. 3B). It should be noted that it is unclear whether αA or αB is sufficient to target GFP to chloroplast membranes when either of them is fused to GFP independent of the CTD.

### Possible mechanisms governing the amphipathic helix-mediated membrane targeting

It is not yet clear how each replication protein is targeted uniquely to a specific organelle membrane. There exist at least two potential nonexclusive mechanisms by which viral proteins target specific membranes: the viral proteins associate with specific lipids enriched on those membranes, or there are host proteins on those membranes that act as receptors for the viral proteins. For cellular amphipathic α-helices that have been extensively studied, some can sense and bind specific lipids, and some are involved in inducing curvatures (Giménez-Andrés *et al*, 2018). For instance, the amphipathic α-helix in yeast transcription repressor Opi1 (overproduction of inositol 1) binds preferentially to phosphatidic acid (PA) over phosphatidylserine (PS) *in vitro* (Hofbauer *et al*, 2018). On the contrary, the amphipathic α-helix in yeast protein Spo2 (sporulation 20) prefers charged lipids and binds PA and PS with similar affinity (Horchani *et al*, 2014). Given that organelle membranes have different lipid compositions (van Meer *et al*, 2008; Zhang *et al*, 2019), it is possible that amphipathic α-helices from BMV 1a, CCMV 1a, and HEV ORF1 may recognize certain lipids present in ER membranes while helix B in CMV 1a and RuV 1a prefer lipids enriched in the vacuole membranes. In the future, the binding properties of these peptides to different organelle membranes can be examined by using *in vitro* assays (Varkey *et al*, 2020) with liposomes composed of different lipids.

We showed that CMV HB-GFP were at both ER and vacuolar membranes, suggesting that CMV HB-GFP was likely transported to the vacuolar membrane first before reaching the ER via the cellular protein secretory pathway. Of note, we have previously found that several components of the cellular COPII (coat protein complex II) pathway were required for BMV 1a’s association with the nER membrane (Li *et al*, 2016), supporting our idea that the protein secretory and protein trafficking pathway is needed for these helix-mediated protein targeting.

In conclusion, we have demonstrated that helix B of BMV 1a is necessary and sufficient for 1a’s targeting to nER membranes, and thus, plays a critical role in initiating BMV VRC formation at the site of viral replication. The conclusion that an amphipathic helix dictates the targeting of a viral replication protein can be extended to several viruses belonging to the alphavirus-like superfamily, revealing a conserved feature among the superfamily encompassing viruses infecting plants, animals, and humans. Given that targeting viral replication proteins to the designated organelle is the initial step to form functional VRCs for genomic replication, a better understanding will provide opportunities to develop antiviral strategies for virus control.

## Materials and Methods

### Yeast strain

The *Saccharomyces cerevisiae* strain YPH500 (*MAT*α *ura3–52, lys2–801, ade2–101, trp1*-Δ*63, his3*-Δ*200, leu2*-Δ*1*) were used in all experiments. Yeast cells were grown at 30 °C in a synthetic defined medium containing 2% raffinose or galactose as the carbon source. Histidine, tryptophan, uracil, leucine, or combinations of them were omitted from the medium to maintain selection for different plasmid combinations. Cells were harvested as described previously (He *et al*, 2019) when the optical density at 600 nm (OD_600_) reached between 0.6 and 1.0. Except for the localization studies and membrane flotation assay for 1a fragments (region E, helix A, helix B, and helix A+B), all other experiments were carried out using His6-tagged BMV1a and WT 2a^pol^.

### Plant Materials and Growth Conditions

All agroinfiltration experiments were performed in WT or H2B-RFP (red fluorescent protein fused to the C terminus of histone 2B) transgenic *N. benthamiana*. Plants were grown in a growth chamber with temperatures at 26°C (16 h, light) and 22°C (8 h, dark) for 4-6 weeks before being infiltrated with agrobacterium cultures. After infiltration, the plants were kept under the same growth conditions.

### Plasmids

BMV 1a, mutant derivatives, and 2a^Pol^ were expressed under the control of the *GAL1* promoter, using pB1YT3-cH6 (with a His6 tag at the C-terminus of BMV 1a) or derivatives, and pB2YT5, respectively (Ahola *et al*, 2000). BMV RNA3 was expressed from pB3MS82, where an RNA3 derivative with an abolished coat protein was under the *GAL1* promoter (Sullivan & Ahlquist, 1999). pB1YT3-mC was used as a vector to incorporate a 32 amino acids linker (-DVGGGGSEGGGSGGPGSGGEGSAGGGSAGGGS-) between 1a and mCherry. pB1YT3-L32-mC or pB1YT3-L32-GFP was used as the vector to introduce mutations in helix B by an overlapping PCR approach using specific primers to introduce mutations and to replace BMV 1a with region E, helix A+B, helix A, or helix B to make pE-, pHAB- pHA-, or pHB-L32-GFP. Organelle markers Scs2-mC, Sec63-GFP, GFP-Sed5 were expressed under the control of an endogenous promoter from a low-copy-number plasmid.

BMV or CMV HB was amplified from yeast plasmid pHB-L32-GFP-BMV or pHB-L32-GFP-CMV and inserted into the *Bam*HI-digested p1300-YFP vector (Zhao *et al*, 2020, 2021) in frame with and upstream of YFP and under the control of the Cauliflower mosaic virus 35S promoter to generate BMV HB-YFP and CMV HB-YFP.

### RNA extraction and northern blotting

For RNA extraction, yeast cells were harvested between OD_600_ 0.6-1.0, and total RNA was extracted by a hot phenol method (Köhrer & Domdey, 1991). Equal amounts of total RNA were used for Northern blot analysis. ^32^P-Labeled probes specific to BMV positive- or negative-strand RNAs or 18S rRNA were used for the hybridization. 18S rRNA was used as a loading control to eliminate loading variations. Radioactive signals of BMV positive-, negative-strand, or 18S rRNA were scanned using a Typhoon FLA 7000 phosphorimaging system, and the intensity of signals was quantified using Adobe Photoshop.

### Protein extraction and western blotting

For protein analysis, two OD_600_ units of yeast cells were harvested and frozen prior to extraction. Total proteins were extracted as described previously (Li *et al*, 2016). Equal volumes of total lysates were analyzed using SDS-PAGE and transferred to a polyvinylidene difluoride (PVDF) membrane (Merck Millipore). Rabbit anti-BMV 1a antiserum (1:10,000 dilution, a gift from Dr. Paul Ahlquist at the University of Wisconsin-Madison), mouse anti-BMV 2a^pol^ (1:3000 dilution, a gift from Dr. Paul Ahlquist at the University of Wisconsin-Madison), mouse anti-Pgk1 (1:10,000 dilution; Thermo Fisher Scientific), anti-GFP (1:5000 dilution), and anti-RFP (1:3000 dilution, GenScript), anti-Dpm1 (1:3,000 dilution, Thermo Fisher Scientific) was used as the primary antibody. Horseradish peroxidase conjugated anti-rabbit or anti-mouse antibody (1:10,000 dilution, Thermo Fisher Scientific) together with the Supersignal West Femto maximum sensitivity substrate (Thermo Fisher Scientific) was used to detect target proteins. Band intensities were quantified using Adobe Photoshop.

### Membrane flotation assays

Flotation assays were performed as described previously (Zhang *et al*, 2012, 2016). Briefly, spheroplasts prepared from ten OD_600_ units of yeast cells expressing His6-tagged helix B point mutants, and deletion mutants 1a-ΔA, and 1a-ΔB, and fragments spanning GFP fused region E, helix A+B, helix A, and helix B were resuspended in 350 μl of TNE buffer (50 mM Tris-HCl [pH 8], 150 mM NaCl, 5 mM EDTA) containing 1:100 dilution of yeast/fungal protease arrest from G biosciences. The resulting cell lysate was centrifuged for 5 min at 500 x *g* to remove cell debris. The supernatant was adjusted to 40% iodixanol by the addition of 60% iodixanol (Axis-Shield, Oslo, Norway). A 600 μl of the mixture was placed at the bottom of a Beckman TLS55 centrifuge tube and overlaid with 1.4 ml of 30% iodixanol in TNE and 100 μl of TNE. The gradients were centrifuged at 201,000 x *g* at 4°C for 5 h. The gradients were divided into 6 fractions (360 μl for the top two fractions and 320 μl for the rest) and analyzed by Western blotting for specific proteins.

### Vacuolar membrane staining

To stain yeast vacuolar membranes, a lipophilic dye, FM4-64 (Thermo Fisher) that selectively stains the vacuolar membranes was used. A stock concentration of 1.6 μM in DMSO was made and staining was performed as described previously (Vida & Emr, 1995). Yeast cells at an OD_600_ of 0.6-0.8 were collected and centrifuged at 5000 x *g* for 5 minutes at RT. The pellet was suspended in 50 μl YAPD and mixed with 1μl of the stock dye. The cells were incubated in a water bath maintained at 30°C for 30 minutes. After the incubation, 1 ml YAPD was added, and the mixture was centrifuged at 5000 x *g* for 5 minutes at RT. The cell pellet was collected and resuspended in 5 ml YAPD and transferred to culture tubes and grown for 2.5 hours. The cells were harvested by centrifugation at 5000 x *g* for 5 minutes and fluorescence was detected at an excitation/emission maximum of ~515/640 nm.

### Structure prediction

RaptorX protein prediction platform was used for obtaining the pdb files for the 1a protein. The solved crystal structure of the tomato mosaic virus helicase-like domain (PDB-3VKW) was automatically chosen as a template by the program to generate the data files. The obtained pdb file was visualized using PyMol.

### Fluorescence and confocal imaging

For all localization studies, yeast cells co-expressing mC- and eGFP-tagged proteins were grown overnight in a synthetic defined medium containing 2% raffinose as a carbon source at 30°C. From this starter culture, a subculture with an OD_600_ of 0.3 was initiated and grown in 2% galactose as a carbon source for the defined hours for microscopic analysis.

For confocal microscopy in plants, the expression vectors p1300-YFP, -BMV HB-YFP, -CMV HB-YFP, and CCMV HB-YFP were individually introduced into *A. tumefaciens* strain GV3101. After overnight growth and activation, Agrobacterium cultures were harvested and resuspended in the induction buffer (10 mM MgSO4, 100 mM 2-N-morpholino ethanesulfonic acid [pH 5.7], 2 mM acetosyringone), and incubated for at least 2 h at room temperature. The suspensions were then adjusted to OD600=1.0 and infiltrated into WT or H2B-RFP transgenic *N. benthamiana* plants. After agroinfiltration, *N. benthamiana* were grown in a growth chamber with a 16 h light/8 h dark cycle. At 40 hpai, leaves were excised and YFP fluorescence was examined in epidermal cells using confocal microscopy (ZEISS LSM 710). The microscope was configured with a 458–515 nm dichroic mirror for dual excitation and a 488-nm beam splitter to help separate YFP fluorescence. Photographic images were prepared using ZEN 2011SP1.

## Acknowledgments

We thank Haijie Liu, Parkesh Suseendran, Xin Zhang, Nicholas Todd and Laily Jaghori for their general help and assistance. We appreciate help from Dr. Xiang-Jin Meng for providing the HEV cDNA fragment and general discussion, and Dr. Kristi DeCourcy at the Fralin Life Science Institute for fluorescence and confocal microscopes.

## Author contributions

Conceptualization: PS, MS, AD, XW

Methodology: PS, WZ, EF, MS, XW

Investigation: PS, WZ, AM, EF, MS, XW

Visualization: PS, WZ, AM, GH, EF

Supervision: MS, XW

Writing—original draft: PS, XW

Writing—review & editing: PS, MS, AD, and XW

## Data and materials availability

All data needed to evaluate the conclusions in the paper are present in the paper and/or the Supplementary Materials. Additional data related to this paper may be requested from the authors.

## Expanded view figures

**Fig. EV1.**
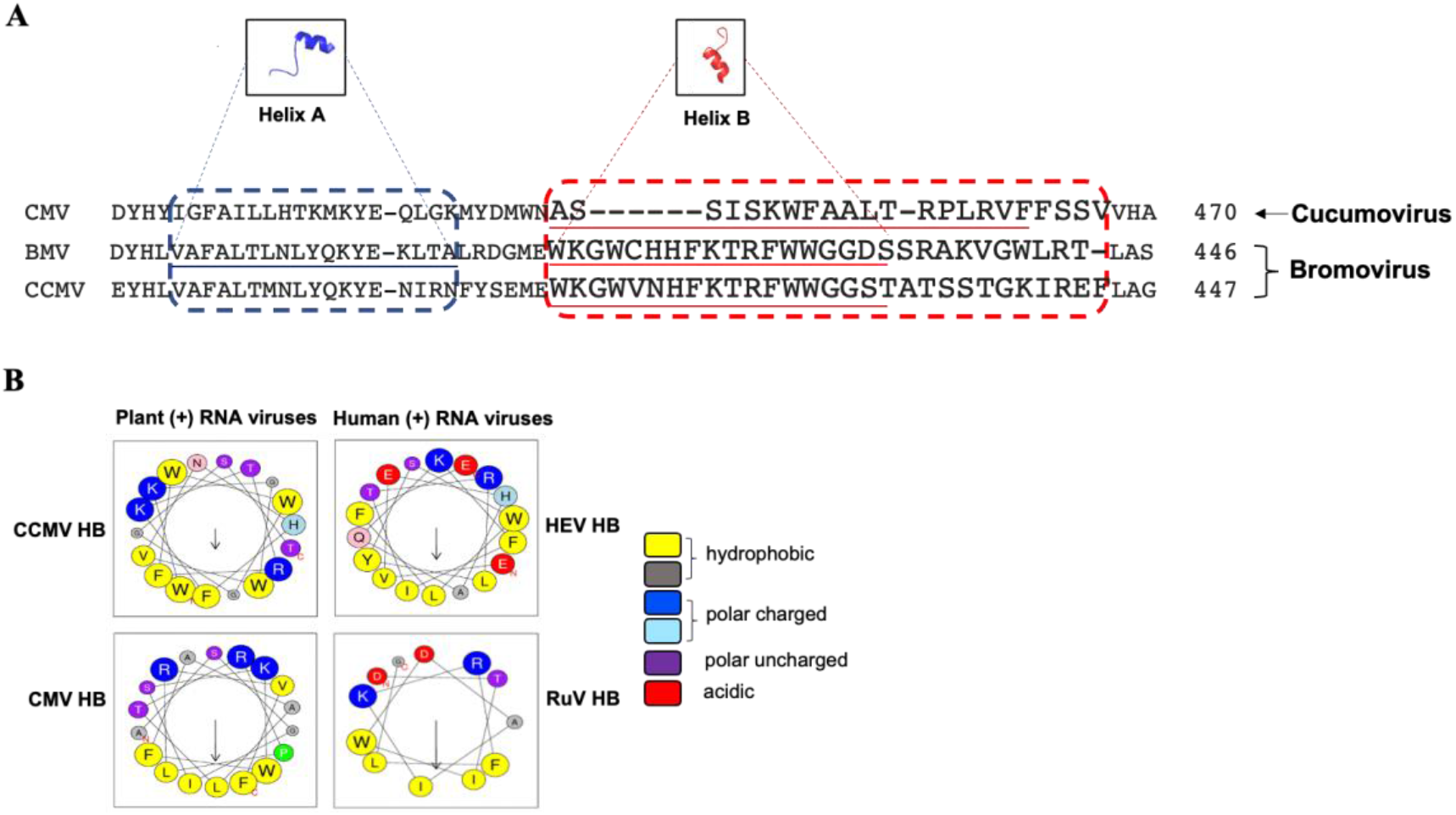
Two amphipathic alpha-helices are present in the replication proteins from members of the alphavirus-like superfamily. **(*A*)** Multiple sequence alignment shown for a short stretch of region encompassing helix A and helix B across plant viruses, BMV, CCMV, and CMV. The members of a genus share higher identity in helix A and helix B (CCMV and BMV) than members from higher ranks such as family (BMV and CMV). **(*B*)** Helical wheel projections of amphipathic helix B of plant (CMV, CCMV) and animal viruses (HEV, RuV) using the predicted helix B regions. Amphipathic faces for CCMV, CMV as predicted by HeliQuest are WGFWFV and PWFLILFA and for HEV and RuV are LALIVY and AFIILW respectively. The arrow in helical wheels corresponds to the hydrophobic moment.

**Fig. EV2.**
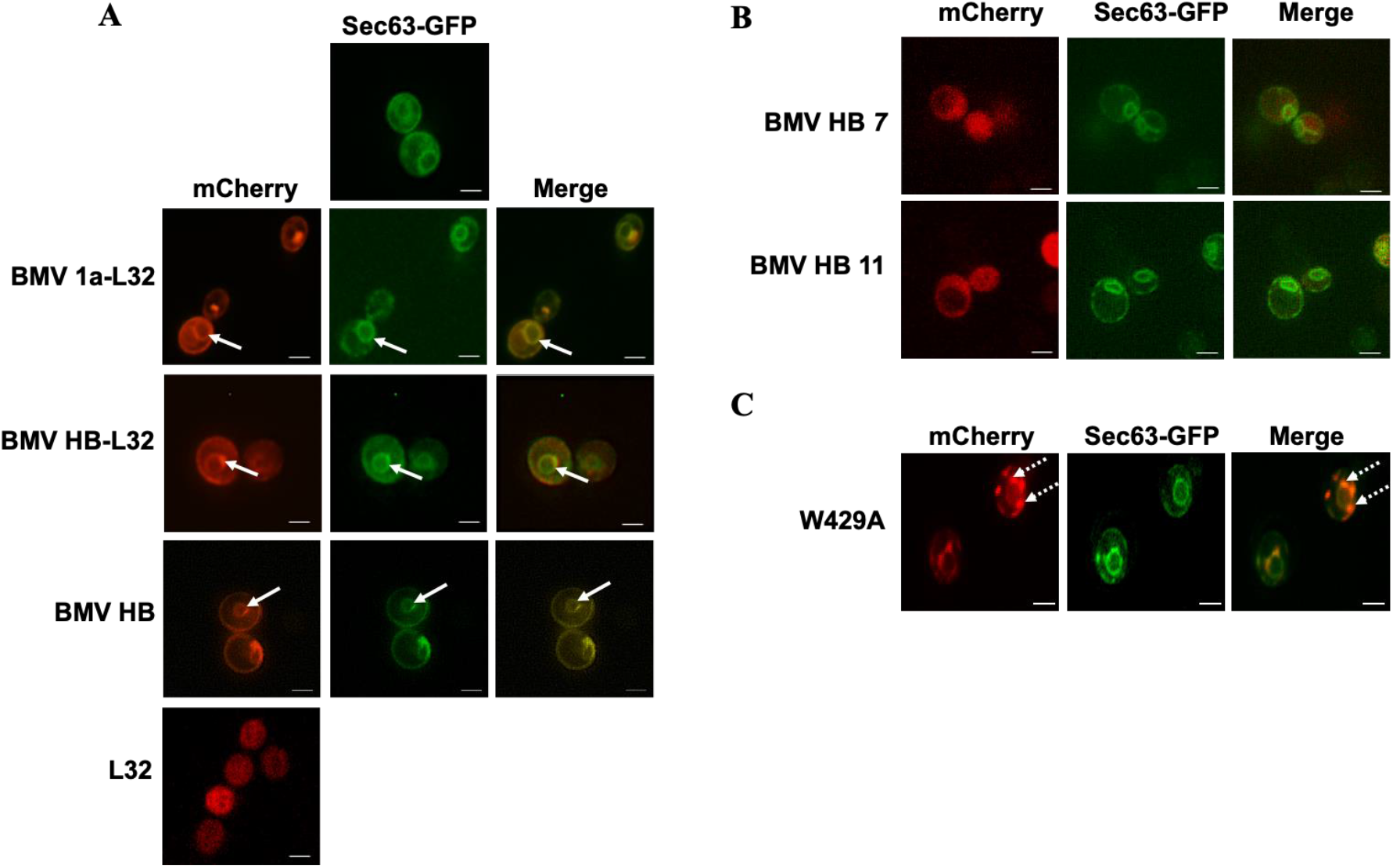
The 32 amino acid-long linker is not required for the correct localization of fluorescent protein-tagged 1a or helix B. **(*A*)** Fluorescence microscopic images showing that mCherry-tagged BMV 1a or helix B with or without the 32 aa-long linker is colocalized with an ER marker Sec63-GFP. L32-mC, which is not fused to BMV 1a or helix B, is localized in the cytosol (bottom panel). **(*B*)** Truncated versions of helix B spanning W416-H422 (BMV HB *7*) or W416-R426 (BMV HB *11*) fail to target mCherry to ER membranes. **(*C*)** Localization of a representative mutant, W429A, that formed puncta similar to mutants F423A and T425A (shown in dotted arrows) in cells co-expressing Sec63-GFP. Scale bars: 2μm.

**Fig. EV3.**
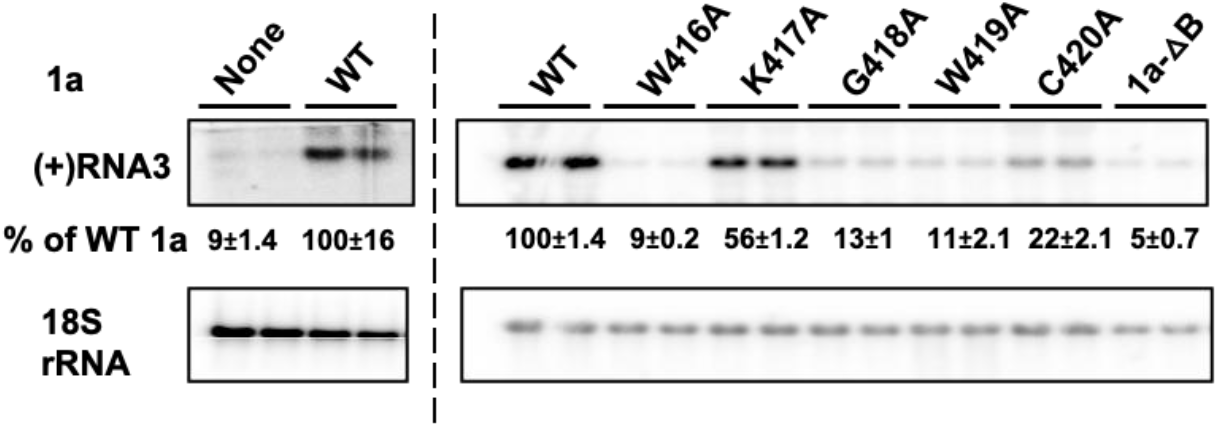
Mutations in BMV 1a helix B inhibit the 1a-mediated RNA3 protection. RNA3 accumulation in the presence of WT or mutant 1a. Total RNA was isolated from cells expressing RNA3 alone or in the presence of WT or mutant 1a. Viral (+)RNA3 was detected using a BMV positive-strand-specific probe. 18S rRNA served as a loading control. Band intensities were measured using Adobe Photoshop. Note Northern blot on the left for WT and the control was done independently from the blot on the right.

**Fig. EV4.**
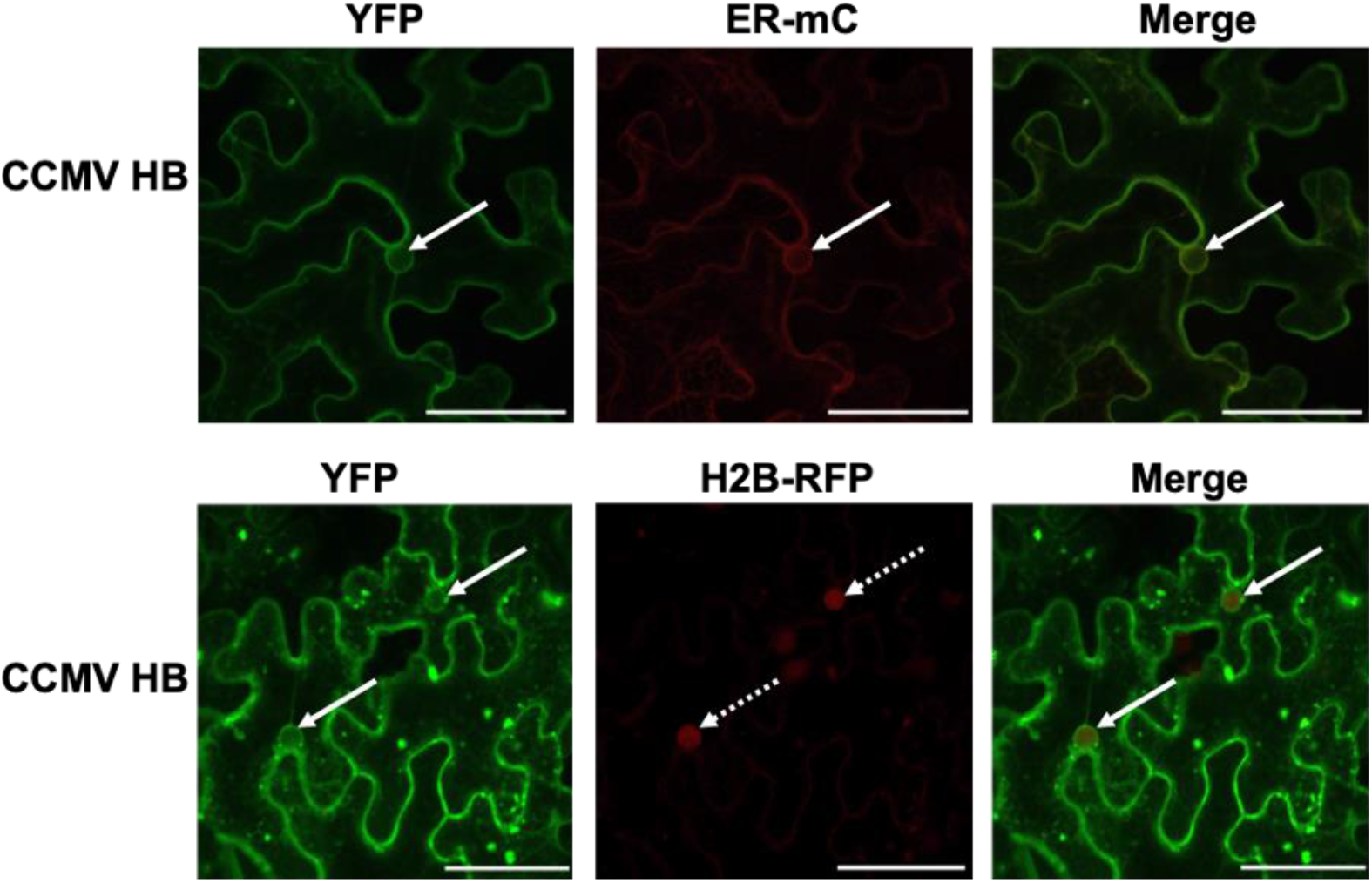
CCMV HB directs yellow fluorescent protein to ER membranes. Confocal microscope images showing YFP-tagged CCMV HB with an ER-mC marker (top row). The bottom panel shows colocalization of YFP-tagged CCMV HB expressed in histone 2B-RFP transgenic plant background. Solid arrows denote the nER localization and dotted arrows represent the nucleus. Scale bars: 50μm.

